# Glycolytic reprogramming underlies immune cell activation by polyethylene wear particles

**DOI:** 10.1101/2022.10.14.512318

**Authors:** Chima V. Maduka, Oluwatosin M. Habeeb, Maxwell M. Kuhnert, Maxwell Hakun, Stuart B. Goodman, Christopher H. Contag

## Abstract

Primary total joint arthroplasties (TJAs) are widely and successfully applied reconstructive procedures to treat end-stage arthritis. Nearly 50% of TJAs are now performed in young patients, posing a new challenge: performing TJAs which last a lifetime. The urgency is justified because subsequent TJAs are costlier and fraught with higher complication rates, not to mention the toll taken on patients and their families. Polyethylene particles, generated by wear at joint articulations, drive aseptic loosening by inciting insidious inflammation associated with surrounding bone loss. Down modulating polyethylene particle-induced inflammation enhances integration of implants to bone (osseointegration), preventing loosening. A promising immunomodulation strategy could leverage immune cell metabolism, however, the role of immunometabolism in polyethylene particle-induced inflammation is unknown. Our findings reveal that immune cells exposed to sterile or contaminated polyethylene particles show fundamentally altered metabolism, resulting in glycolytic reprogramming. Inhibiting glycolysis controlled inflammation, inducing a pro-regenerative phenotype that could enhance osseointegration.

## Introduction

End-stage arthritis can be successfully treated by primary total joint arthroplasties (TJAs)^1^. With nearly 50% of TJAs performed in patients younger than 65 years^2^, the vision of TJAs is now to reconstruct joints which will last a lifetime, despite patients’ daily activities^3^. This is especially crucial because revision TJAs are costlier and fraught with higher complication rates, technical difficulties, and poorer surgical outcomes than primary TJAs^4^. Such revision TJAs commonly arise from aseptic loosening, frequently incited by polyethylene wear particles generated by relative motion at joint articulations^5^. Aseptic loosening may occur with or without adsorbed contaminants, such as bacterial lipopolysaccharides (LPS). Wear particles induce prolonged, low-grade inflammation with macrophages and fibroblasts as key immune cellular players^6^. This pathology is often radiographically detected only when surrounding bone loss (periprosthetic osteolysis) occurs^3^. By then, compromised implant stability results in loosening and implant failure, necessitating revision surgeries.

To minimize generation of wear particles, ultrahigh molecular weight polyethylene liners at the bearing surfaces of reconstructed joints are currently being replaced by highly crosslinked polyethylene. Crosslinked polyethylene has significantly reduced the amount of generated wear particles and accompanied chronic inflammation with periprosthetic osteolysis^7^. However, crosslinking does not completely block the generation of wear particles from bearing surfaces of implants and subsequent inflammation^8^. Up to 9% of patients with crosslinked polyethylene liners present with chronic inflammation-induced periprosthetic osteolysis 15 years later^9^. Moreover, crosslinking has little effect on particles from third body wear, backside wear and impingement^10^; and there are currently no agents that specifically treat polyethylene particle-induced inflammatory osteolysis^11^. Consequently, there is an unmet clinical need to develop methods that will mitigate aseptic loosening from polyethylene particle-induced chronic inflammation to improve implant longevity. Furthermore, as particles generated from ultrahigh molecular weight or highly crosslinked polyethylene similarly result in inflammation^8,11^, either of them effectively models particle-induced inflammation.

Metabolic reprogramming refers to changes in glycolytic flux and oxidative phosphorylation (OXPHOS), traditional bioenergetic pathways, that are inextricably linked to macrophage activation toward proinflammatory^12,13^ or pro-regenerative phenotypes^14,15^. Advances in understanding macrophage-mesenchymal stem cell crosstalk^16^ has revealed that down modulating inflammation induced by polyethylene particles can prevent implant loosening by enhancing osseointegration through increased pro-regenerative macrophage activity. For example, using mesenchymal stem cells (MSCs)^17^ and engineered IL-4 expressing MSCs^18^; targeting inflammatory pathways using decoy molecules for NF-kB^19^, TNF-a^20^ and MCP-1^21^; and using antioxidants like vitamin E^11^ have shown promise for enhanced osseointegration by reducing inflammation. However, the metabolic underpinnings underlying macrophage activation by polyethylene particles are largely undefined. A detailed understanding of metabolic programs could be leveraged for immunomodulation toward extending the longevity of implants. Here, we show that both macrophages and fibroblasts exposed to sterile or LPS-contaminated polyethylene particles undergo metabolic reprogramming and differential changes in bioenergetics. Glycolytic reprogramming underlies increased levels of proinflammatory cytokines, including MCP-1, IL-6, IL-1β and TNF-a. Specific inhibition of different glycolytic steps not only modulated these proinflammatory cytokines but stimulated pro-regenerative cytokines, including IL-4 and IL-10, without affecting cell viability. Concomitant elevation of both glycolytic flux and oxidative phosphorylation by polyethylene particles and inhibitory effects on inflammatory cytokines in addition to IL-1β^13^ suggest a unique metabolic program that could be targeted for pro-regenerative clinical outcomes following TJAs.

## Results

### Bioenergetics is differentially altered in immune cells exposed to polyethylene particles

We had previously optimized an in-vitro, live-cell, bioenergetic workflow where ATP is rate-limiting to measure spatiotemporal bioenergetic alterations in cells exposed to biomaterials^22^. This involved transfecting mouse embryonic fibroblasts (MEFs) with a Sleeping Beauty transposon plasmid (pLuBIG) having a bidirectional promoter driving an improved firefly luciferase gene (fLuc) and a fusion gene encoding a Blasticidin-resistance marker (BsdR) linked to eGFP (BGL)^23^. Both highly crosslinked^8^ and ultrahigh molecular weight^21^ polyethylene particles similarly incite inflammation and are clinically used. Ultrahigh molecular weight polyethylene particles whose doses and sizes have been previously characterized were examined herein after polyethylene particles were determined to be endtoxin-free^17–19,21^. Since adsorbed bacterial lipopolysaccharide (LPS; a.k.a. endotoxin) could play a role in aseptic loosening^24^, we compared key results to cells exposed to polyethylene particles and LPS.

Whereas only polyethylene particles consistently lowered bioenergetic (ATP) levels in live BGL cells, overall, LPS alone did not affect ATP levels when compared to untreated fibroblasts over time (Fig. 1a). In comparison to polyethylene particles or LPS alone, combining polyethylene particles and LPS further decreased ATP levels after prolonged exposure (Fig. 1a). D-luciferin used in live-cell assays could be limited by its ability to permeate cell membranes^25^; accordingly, bioenergetic measurement in lysed fibroblasts was more sensitive, corroborating decreases in ATP levels after exposure to only polyethylene particles (by 1.2-fold) or a combination of polyethylene particles and LPS (by 1.1-fold) relative to untreated cells at day 3 (Fig. 1b). Primary bone marrow-derived macrophages revealed a 1.5-, 1.8-, and 1.6-fold decrease in ATP levels relative to untreated cells following exposure to only polyethylene particles, only LPS, and polyethylene particles with LPS, respectively (Fig. 1c).

**Figure 1.**
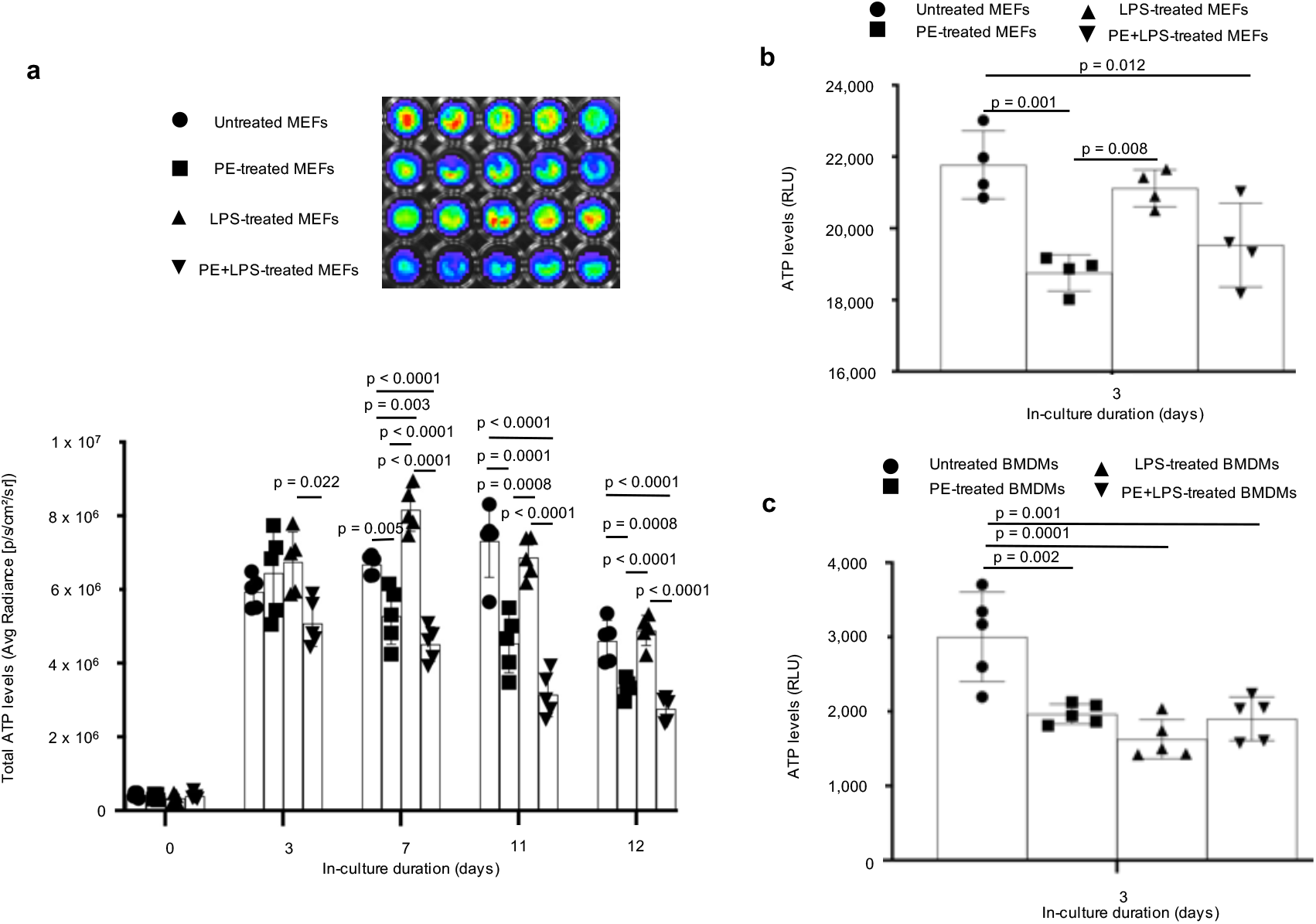
Ultrahigh molecular weight polyethylene (PE) particles, alone or in combination with endotoxin (LPS), alter bioenergetic (ATP) levels/. **a**, Over time, PE particles lower bioenergetics in blasticidin-eGFP-luciferase (BGL)-transfected mouse embryonic fibroblasts (MEFs) compared to untreated cells; combining PE particles and LPS lowers ATP levels compared to PE particles or LPS alone (representative bioluminescent wells shown). **b**, In lysed wild-type MEFs, bioenergetics is lowered after exposure to PE particles. **c**, In primary bone marrow-derived macrophages (BMDMs), PE particles and LPS, alone or in combination, decrease bioenergetics. Mean (SD), n = 5 (Fig. 1a, 1c), n = 4 (Fig. 1b) one-way ANOVA followed by Tukey’s post-hoc test.

### Exposure to polyethylene particles alters functional metabolism in immune cells

To explore what bioenergetic pathways were responsible for alterations in ATP levels, we used the Seahorse assay to probe extracellular acidification rate (ECAR), lactate-linked proton efflux rate (PER) and oxygen consumption rate (OCR). ECAR, PER and OCR are indices of glycolytic flux, monocarboxylate transporter (MCT) function^26,27^ and mitochondrial oxidative phosphorylation, respectively, and are used to assess metabolic reprogramming^12,13^. Following exposure to LPS alone, fibroblasts did not reveal changes in ECAR, PER or OCR compared to untreated cells (Fig. 2a-c). In contrast, exposure to polyethylene particles resulted in a 1.7-, 1.7-, and 2-fold increase in ECAR, PER and OCR, respectively, relative to untreated fibroblasts (Fig. 2a-c). Similarly, a combination of polyethylene particles and LPS increased OCR by 1.6-fold in comparison to untreated fibroblasts (Fig. 2c).

**Figure 2.**
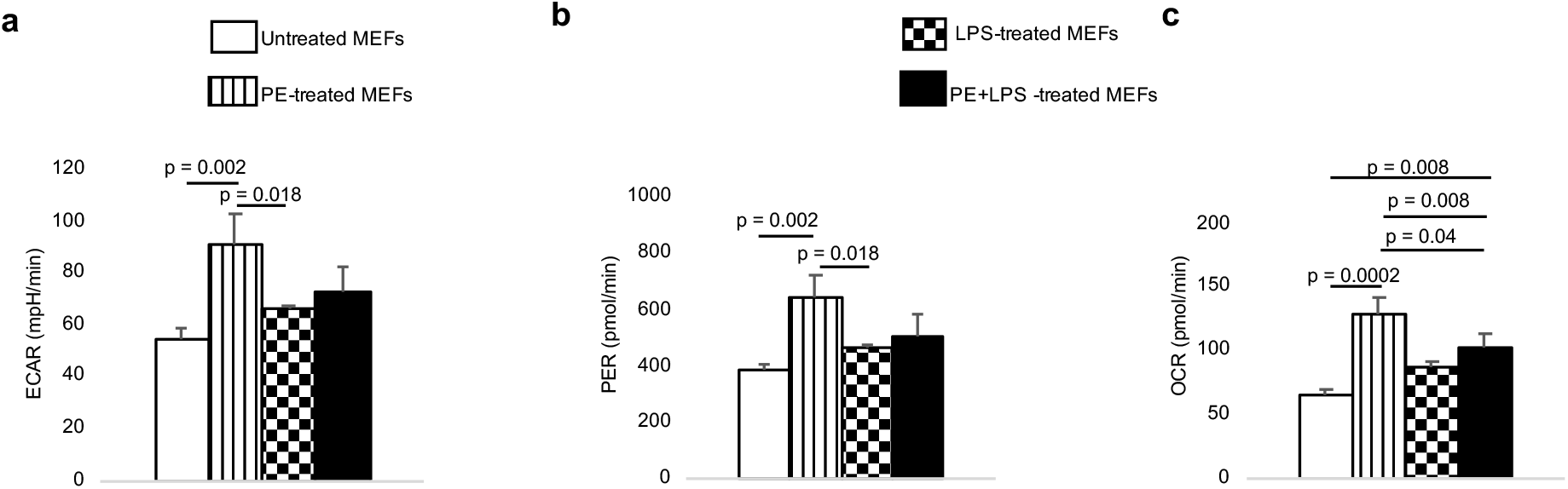
Mouse embryonic fibroblasts (MEFs) exposed to ultrahigh molecular weight polyethylene (PE) particles alone show increased functional metabolic indices. **a-c**, In comparison to untreated cells, PE particle-treated MEFs have higher extracellular acidification rate (ECAR; **a**), proton efflux rate (PER; **b**) and oxygen consumption rate (OCR; **c**). Mean (SD), n = 3, one-way ANOVA followed by Tukey’s post-hoc test.

Exposure to only polyethylene particles increased ECAR, PER and OCR by 13.1-, 13.1- and 3.1-fold, respectively, in primary macrophages compared to untreated cells (Fig. 3a, c, e). Macrophages exposed to polyethylene particles and LPS increased ECAR, PER and OCR by 23-, 23.1- and 2.8-fold, respectively, compared to untreated cells (Fig. 3b, d, f).

**Figure 3.**
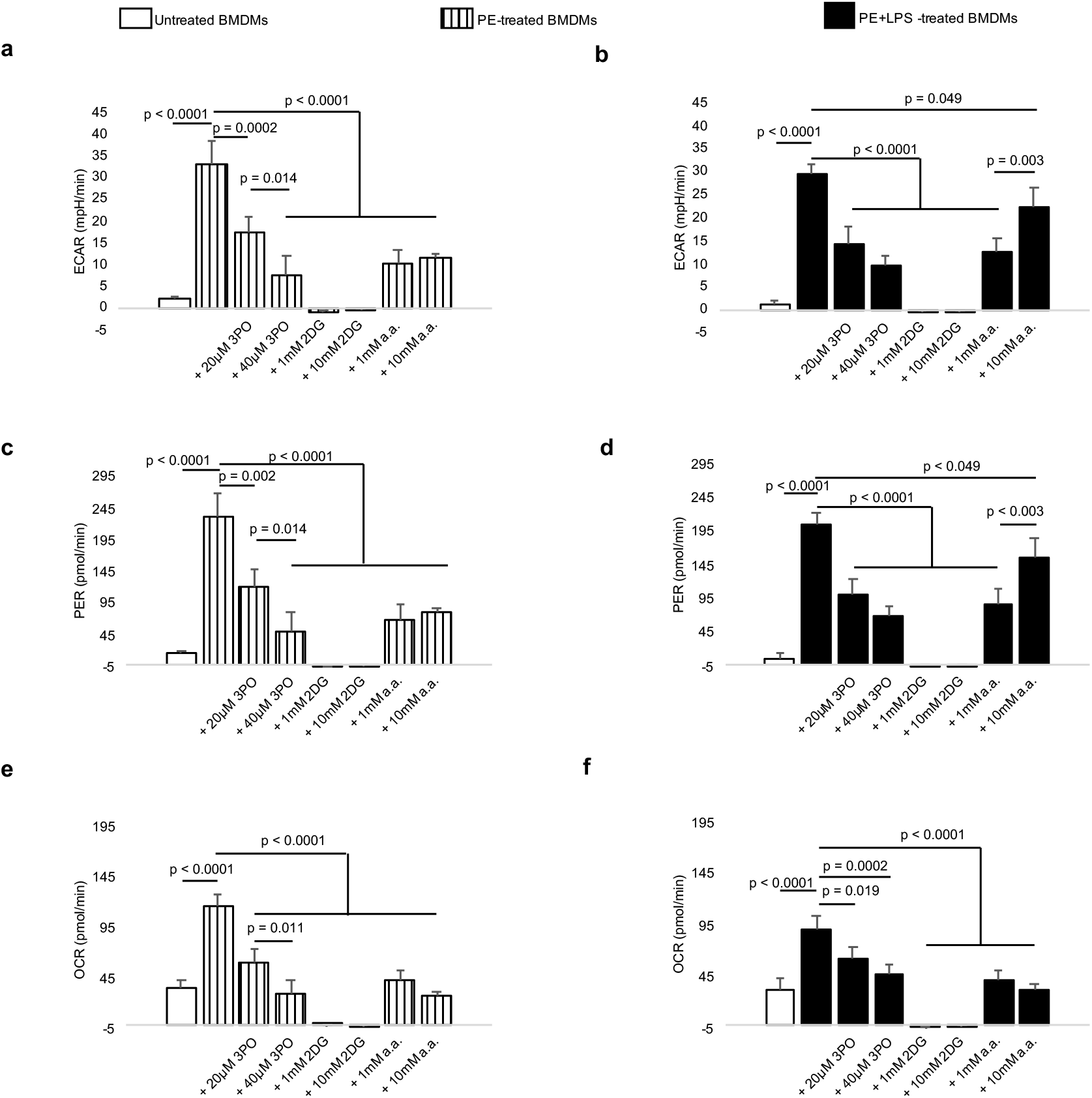
Primary bone marrow-derived macrophages (BMDMs) exposed to ultrahigh molecular weight polyethylene (PE) particles or both PE particles and endotoxin (LPS) reveal greater extracellular acidification rate (ECAR), proton efflux rate (PER) and oxygen consumption rate (OCR) than untreated cells; this increment is reduced upon addition of various glycolytic inhibitors. **a-f,** ECAR (**a-b**), PER (**c-d**) and OCR (**e-f**) are increased in BMDMs treated with PE particles, alone or in combination with LPS; elevated levels are decreased upon addition of 3-(3-Pyridinyl)-1-(4-pyridinyl)-2-propen-1-one (3PO), 2-deoxyglucose (2DG) or aminooxyacetic acid (a.a.). Mean (SD), n = 3, one-way ANOVA followed by Tukey’s post-hoc test.

To reduce abnormal increments in ECAR, PER and OCR, we targeted different stages of glycolysis using 3-(3-pyridinyl)-1-(4-pyridinyl)-2-propen-1-one (3PO), 2-deoxyglucose (2DG) and aminooxyacetic acid (a.a.). 3PO inhibits 6-phosphofructo-2-kinase which is the rate limiting glycolytic enzyme^28^; 2DG inhibits hexokinase, the first enzyme in glycolysis^13^; and a.a. prevents the mitochondrion from utilizing glycolytic pyruvate^29^. In a dose-dependent manner, 3PO, 2DG and a.a. decreased ECAR, PER and OCR among macrophages exposed to only polyethylene particles or a combination of polyethylene particles and LPS (Fig. 3a-f), suggesting efficient cellular uptake and pharmacologic effects of these small molecule inhibitors.

Compared to untreated cells, there was no difference in cell numbers following exposure to polyethylene particles, LPS or polyethylene particles with LPS among fibroblasts (Fig. 4a) or macrophages (Fig. 4b). Additionally, exposure of macrophages to pharmacologic inhibitors, including 3PO, 2DG and a.a. did not lower cell viability (Fig. 4c). Importantly, in fibroblasts exposed to polyethylene particles alone or polyethylene particles and LPS, addition of 3PO, 2DG or a.a. further lowered bioenergetics in a dosedependent manner (Fig. 5).

**Figure 4.**
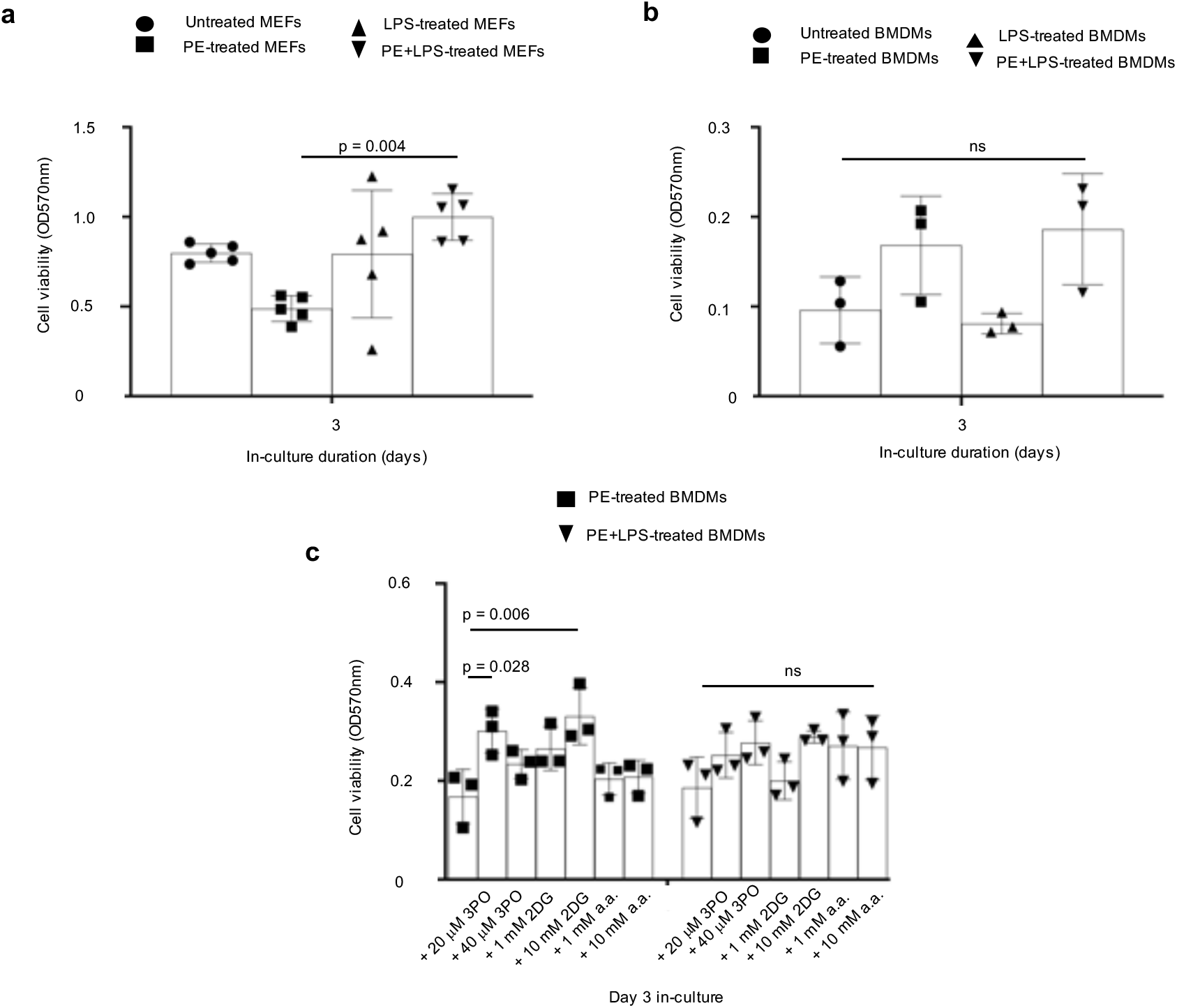
Compared to untreated cells, treatment with ultrahigh molecular weight polyethylene (PE) particles, endotoxin (LPS) or a combination of PE particles and LPS does not change cell numbers; addition of glycolytic inhibitors does not decrease cell numbers. **a-b**, In mouse embryonic fibroblasts (MEFs; **a**) or primary bone marrow-derived macrophages (BMDMs; **b**), exposure to PE particles, LPS or PE particles and LPS does not change cell numbers relative to untreated controls. **c**, Addition of various doses of 3-(3-Pyridinyl)-1-(4-pyridinyl)-2-propen-1-one (3PO), 2-deoxyglucose (2DG) or aminooxyacetic acid (a.a.) to PE particle-treated or PE particle- and LPS-treated BMDMs does not decrease cell numbers. Mean (SD), n = 5 (Fig. 4a), n = 3 (Fig. 4b, c), one-way ANOVA followed by Tukey’s post-hoc test.

**Figure 5.**
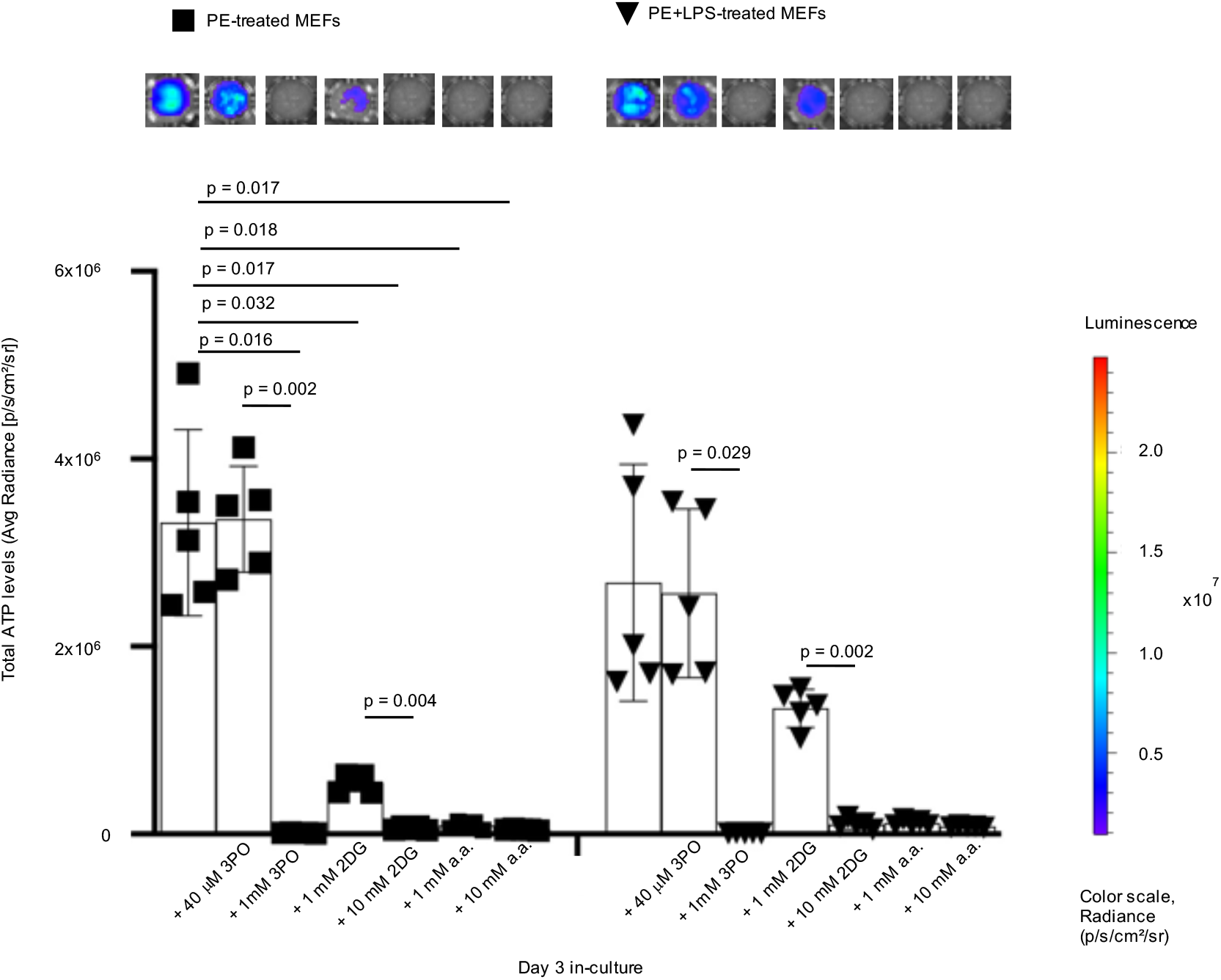
Glycolytic inhibitors decrease bioenergetic levels in treated mouse embryonic fibroblasts (MEFs). Following treatment of blasticidin-GFP-Luciferase (BGL)-transfected MEFs with ultrahigh molecular weight polyethylene (PE) particles alone or in combination with endotoxin (LPS), addition of 3-(3-pyridinyl)-1-(4-pyridinyl)-2-propen-1-one (3PO), 2-deoxyglucose (2DG) and aminooxyacetic acid (a.a.; representative wells are shown) tend to decrease bioenergetics in a dose-dependent manner. Not significant (ns), mean (SD), Brown-Forsythe and Welch ANOVA followed by Dunnett’s T3 multiple comparisons test, n = 5.

### Immunometabolism underlies macrophage polarization by polyethylene particles

To evaluate how metabolism affects immune cellular function, we assayed levels of cytokine and chemokine expression using a magnetic bead-based technique^30^. We observed that proinflammatory proteins, including MCP-1 (Fig. 6a), IL-6 (Fig. 6b), IL-1β (Fig. 6c) and TNF-a (Fig. 6d) were increased by 4.1-, 97.3-, 41.8- and 7-fold, respectively, after exposure to polyethylene particles in comparison to untreated macrophages.

**Figure 6.**
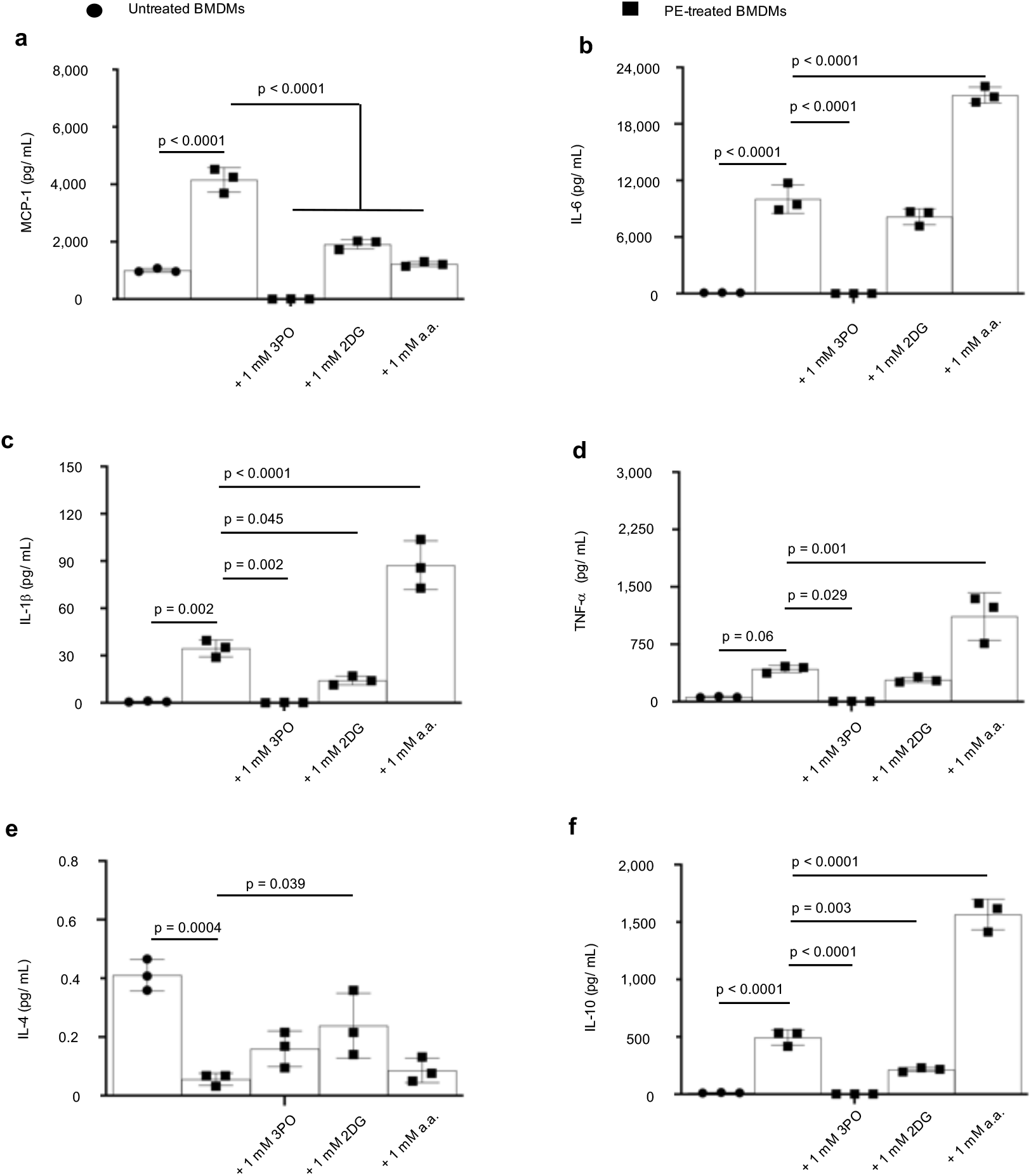
Elevated proinflammatory cytokine (protein) levels are decreased following addition of glycolytic inhibitors to primary bone marrow-derived macrophages (BMDMs). **a-d**, In BMDMs, exposure to ultrahigh molecular weight polyethylene (PE) particles increase proinflammatory cytokines, including MCP-1 (**a**), IL-6 (**b**), IL-1β (**c**) and TNF-a (**d**) in comparison to untreated BMDMs. Addition of 3-(3-Pyridinyl)-1- (4-pyridinyl)-2-propen-1-one (3PO) or 2-deoxyglucose (2DG) decreases proinflammatory cytokines; aminooxyacetic acid (a.a.) selectively decreases MCP-1 levels. **e**, Exposure of BMDMs to PE particles decreases IL-4 levels in comparison to untreated cells; IL-4 levels tend to increase following addition of glycolytic inhibitors. **f**, Compared to BMDMs exposed to only PE particles, exposure to PE particles and a.a. increase IL-10 levels. Mean (SD), n = 3, one-way ANOVA followed by Tukey’s post-hoc test; assay was performed after 3 days in-culture.

Addition of 3PO or 2DG consistently decreased proinflammatory cytokine or chemokine levels (Fig. 6a-d) relative to macrophages exposed to only polyethylene particles; however, addition of a.a. selectively decreased MCP-1 expression (Fig. 6a). Exposure of macrophages to polyethylene particles decreased IL-4 levels by 7.4-fold compared to untreated cells; addition of 3PO, 2DG or a.a. increased IL-4 levels by 2.9-, 4.3-, and 1.5-fold, respectively, relative to polyethylene particles alone; however, only the increase by 2DG was statistically significant (Fig. 6e). Levels of IL-13 and IFN-λ were unchanged (data not shown). Consistent with macrophage polarization being a continuum^31,32^, polyethylene particles increased IL-10 expression in comparison to untreated macrophages (Fig. 6f). Whereas addition of 3PO or 2DG did not increase IL-10 levels, a.a. increased IL-10 expression by 3.2-fold relative to macrophages exposed to only polyethylene particles (Fig. 6f).

## Discussion

When macrophages are exposed to bacterial lipopolysaccharide (LPS), their bioenergetic (ATP) levels are decreased as part of cell activation and inflammation^33^. This results from reprogrammed metabolism that shifts bioenergetic dependence from mitochondrial oxidative phosphorylation (OXPHOS) to glycolysis, with crucial consequences on proinflammatory^12,13^ and anti-inflammatory^14,15^ events. While immunometabolism in response to LPS has been well characterized for such clinical applications as bacterial sepsis, the role of immunometabolism in sterile inflammation induced by clinically relevant implant materials is unknown.

Macrophages are the dominant immune cell type implicated in the chronic inflammatory response to ultrahigh molecular weight polyethylene (PE) particles^2^, likely acting through Toll-like receptors (TLRs)^34,35^. Following exposure to PE particles of particular sizes and over a threshold, transcriptional signaling occurs through NF-kB^36^, MyD88^37^ and chemerin/ChemR23^38^. Consequently, there is increased production of proinflammatory cytokines that accompany resulting pathologies, including periprosthetic osteolysis. Likewise, fibroblasts play a synergistic role with macrophages. Fibroblasts exposed to PE particles ^39,40^ express MCP-1, RANKL, IL-1β, IL-6, MMP1 and MMP2 which activate osteoclasts, accentuate inflammation and degrade surrounding bone extracellular matrix.

Adsorbed LPS could be a contaminant on sterilized implants and has been documented in a subset of patients diagnosed with aseptic loosening of implants from chronic inflammation^24^, and could exacerbate PE particle-induced inflammation^41^. Therefore, PE and LPS-contaminated PE (cPE) particles were examined and compared to LPS. Our findings reveal that bioenergetic imbalances differentially occur in macrophages and fibroblasts exposed to PE particles, LPS or cPE particles. For example, although LPS did not affect ATP levels in fibroblasts, PE particles lowered cellular bioenergetics. Furthermore, fibroblasts exposed to PE particles but not LPS were metabolically reprogrammed, revealing increases in glycolysis, OXPHOS and monocarboxylate transporter (MCT) function. On the other hand, decreased ATP levels were observed in primary bone marrow-derived macrophages exposed to PE particles, LPS or cPE particles consistent with reliance on glycolysis. Immune cells depend on glycolysis during inflammatory activation as glycolysis produces ATP quicker than OXPHOS, albeit OXPHOS results in overall higher ATP levels. Additionally, this switch to glycolysis is crucial for IL-1β production by stabilizing HIF-1a in macrophages^13^ and fibroblast activation in fibrosis^42^. Surprisingly, in addition to elevated glycolysis, OXPHOS was increased in macrophages exposed to PE or cPE particles, independent of changing cell numbers. Concomitant elevation in both glycolysis and OXPHOS suggests a unique metabolic reprogram induced by PE particles relative to LPS; LPS increases glycolysis while reducing OXPHOS^12^. Accompanied decrease in ATP levels suggests that increased OXPHOS is directed at functions other than cellular energy supply. In a septic model, LPS was shown to repurpose mitochondrial function toward superoxide formation in macrophages^12^. At earlier time points than used in this study, LPS decreased OXPHOS^12^, likely reflecting as yet uncharacterized temporal changes in metabolic reprogramming. Notably, glycolytic flux and MCT function but not OXPHOS were higher in macrophages exposed to cPE than PE particles, relative to respective controls. This may likely be from synergistic signaling with cPE particles, as PE particles and LPS are known to activate TLR2 and TLR4 receptors, respectively^34,35^.

Elevated glycolytic flux in macrophages exposed to PE or cPE particles could be lowered by specific pharmacologic inhibition of different glycolytic steps using 3-(3- pyridinyl)-1-(4-pyridinyl)-2-propen-1-one (3PO)^28^, 2-deoxyglucose (2DG)^13^ and aminooxyacetic acid (a.a.)^29^. Lactate from glycolysis is converted to pyruvate which feeds mitochondrial OXPHOS, and proton-linked lactate is bidirectionally shuttled through MCT^26,27^. Consequently, pharmacologic inhibition of glycolysis lowered aberrantly elevated OXPHOS and MCT function. Pharmacologic inhibition did not result in reduced cell viability, excluding potential toxicity. Using fibroblasts expressing luciferase, we observed that glycolytic inhibition further reduced ATP levels following exposure to PE particles, corroborating cellular bioenergetic dependence on glycolysis.

Contrasting non-degradable PE, polylactide (PLA) is a biodegradable biomaterial. Degradation products of PLA, oligomers and monomers of lactic acid, increase ATP levels only after prolonged exposure to immune cells, modeling in-vivo conditions^22^. Increased ATP levels are the result of elevated flux in both glycolysis and mitochondrial respiration, in-vitro. Corroborating in-vitro observations, radiolabeled glucose uptake was elevated in a subcutaneous model and shown to drive inflammation to sterile PLA; targeting glycolysis in-vivo decreased inflammatory markers, including CD86, by reducing radiolabeled glucose uptake. Similarly, in patients who have undergone total joint arthroplasties, chronic inflammation by PE particles is often diagnosed by increased glycolytic flux. Glycolytic flux is measured using fluorodeoxyglucose in positron emission tomography (PET) combined with computed tomography (CT) or magnetic resonance imaging (MRI)^43–46^. Our findings suggests that PET imaging is enabled by glycolytic reprogramming of immune cells in inflamed joints, following exposure to PE or cPE particles.

Macrophages exposed to PE particles became polarized to a proinflammatory phenotype as measured by elevated protein expression of MCP-1, IL-6, IL-1β and TNF- a. Additionally, IL-10 was increased, consistent with macrophage polarization being a spectrum^31^. Both IL-1β and TNF-a induce RANKL expression which drives osteoclast maturation and differentiation, together with M-CSF^6^. Osteolysis, associated with PE particle-induced chronic inflammation, is the result of net bone loss from osteoclast-mediated bone resorption exceeding osteoblast-mediated bone formation. Similarly, IL-6^47^ and MCP-1^48^ are associated with increased osteolysis and cartilage destruction. Interestingly, 2DG and 3PO decreased aberrantly elevated proinflammatory cytokines. In particular, 2DG allowed for some level of proinflammatory cytokine expression. This is clinically important because a suitable level of inflammation is required for tissue repair and osseointegration^49^; compromised osseointegration is a leading cause of implant failure^4^. Remarkably, whereas 2DG decreased MCP-1, IL-6, IL-1β and TNF-a protein levels which were elevated by PE particles, 2DG is known to selectively decrease IL-1β protein levels from LPS^13^, suggesting unique differences. In contrast to 2DG and 3PO, a.a. selectively decreased MCP-1 but not IL-6, IL-1β and TNF-a; and increased IL-10 levels. Central to macrophage-stem cell crosstalk, IL-10 signaling is critical for tissue regeneration^50^. Glycolytic inhibition using 2DG increased IL-4 levels which were reduced by PE particles. Increment of IL-4 levels suggest a pro-regenerative macrophage phenotype. Acute and chronic inflammation as well as bone loss induced by PE particles is reversed by inducing a pro-regenerative macrophage phenotype using IL-4^18^.

In conclusion, all clinically relevant biomaterials undergo wear at articulations, resulting in different levels of chronic inflammation and undermining the longevity of biomaterials used in arthroplasties. By characterizing immune cell metabolism as being pivotal in the inflammatory pathology induced by polyethylene particles, we reveal a unique vulnerability which could be harnessed for the dual purposes of controlling inflammation and stimulating pro-regenerative immune cell phenotypes. Targeting immunometabolism can be extended to other implant materials^51,52^, improving osseointegration and long-term clinical outcomes for patients undergoing various arthroplasties.

## Methods

### Materials

Ultrahigh molecular weight polyethylene particles were sourced, characterized and determined to be endotoxin-free as previously described^18^. Concentrations of 100ng/ mL of lipopolysaccharide (LPS) from *Escherichia coli* O111:B4 (MilliporeSigma) and 1.25 mg/ mL of ultrahigh molecular weight polyethylene particles were used. Furthermore, 3- (3-pyridinyl)-1-(4-pyridinyl)-2-propen-1-one (MilliporeSigma), 2-deoxyglucose (MilliporeSigma) and aminooxyacetic acid (Sigma-Aldrich) were used for glycolytic inhibition.

### Bioenergetic measurement

Bioluminescence was measured using the IVIS Spectrum in vivo imaging system (PerkinElmer) after adding 150 μg/mL of D-luciferin (PerkinElmer). Living Image (Version 4.5.2, PerkinElmer) was used for acquiring bioluminescence on the IVIS Spectrum. Standard ATP/ADP kits (Sigma-Aldrich) containing D-luciferin, luciferase and cell lysis buffer were used to according to manufacturer’s instructions. Luminescence at integration time of 1,000 ms was obtained using the SpectraMax M3 Spectrophotometer (Molecular Devices) using SoftMax Pro (Version 7.0.2, Molecular Devices).

### Cells

Mouse embryonic fibroblast (MEFs) cell line (NIH 3T3 cell line; ATCC) and primary bone-marrow derived macrophages (BMDMs) derived from C57BL/6J mice (Jackson Laboratories) of 3-4 months^12,53^ were used. NIH 3T3 cells were stably transfected with a Sleeping Beauty transposon plasmid (pLuBIG) having a bidirectional promoter driving an improved firefly luciferase gene (fLuc) and a fusion gene encoding a Blasticidin-resistance marker (BsdR) linked to eGFP (BGL)^23^. This enabled us to monitor bioenergetic changes in live cells^22^. For temporal (IVIS) experiments lasting 12 days, 5,000 BGL cells were initially seeded in each well of a 96-well tissue culture plate in 200 μL of complete medium (see below). For ATP, crystal violet and Seahorse assays, 20,000 wild-type MEFs were seeded. For ATP, crystal violet and cytokine/ chemokine assays, 50,000 BMDMs were seeded; 60,000 BMDMs were seeded for Seahorse experiments. For IVIS experiments with glycolytic inhibitors, 20,000 BGL cells were initially seeded. All time points are indicated on respective graphs. Complete medium comprised of DMEM medium, 10% heat-inactivated Fetal Bovine Serum and 100 U/mL penicillin-streptomycin (all from ThermoFisher Scientific).

### Cell viability

Cell viability was assessed using the crystal violet assay^54^. Absorbance (optical density) was acquired at 570 nm using the the SpectraMax M3 Spectrophotometer (Molecular Devices) and SoftMax Pro software (Version 7.0.2, Molecular Devices).

### Functional metabolism

Basal measurements of oxygen consumption rate (OCR), extracellular acidification rate (ECAR) and lactate-linked proton efflux rate (PER) were obtained in real-time using the Seahorse XFe-96 Extracellular Flux Analyzer (Agilent Technologies)^12,13,15^. Prior to running the assay, cell culture medium was replaced by the Seahorse XF DMEM medium (pH 7.4) supplemented with 25 mM D-glucose and 4 mM Glutamine. The Seahorse ATP rate assay was run according to manufacturer’s instruction and all reagents for the Seahorse assays were sourced from Agilent Technologies. Wave software (Version 2.6.1) was used to export Seahorse data directly as means ± standard deviation (SD).

### Chemokine and cytokine measurements

Cytokine and chemokine levels were measured using a MILLIPLEX MAP mouse magnetic bead multiplex kit (MilliporeSigma)^30^ to assess for IL-6, MCP-1, TNF-a, IL-1β, IL-4, IL-10, IFN-λ and 1L-13 protein expression in supernatants. Data was acquired using Luminex 200 (Luminex Corporation) by the xPONENT software (Version 3.1, Luminex Corporation). Using the glycolytic inhibitor, 3PO, expectedly decreased cytokine values to < 3.2 pg/ mL in some experiments. For statistical analyses, those values were expressed as 3.1 pg/ mL. Values exceeding the dynamic range of the assay, in accordance with manufacturer’s instruction, were excluded. Additionally, IL-6 ELISA kits (RayBiotech) for supernatants were used according to manufacturer’s instructions.

### Statistics and reproducibility

Statistical software (GraphPad Prism) was used to analyse data presented as mean with standard deviation (SD). Significance level was set at p < 0.05, and details of statistical tests and sample sizes, which are biological replicates, are provided in figure legends. Exported data (mean, SD) from Wave in Seahorse experiments had the underlying assumption of normality and similar variance, and thus were tested using corresponding parametric tests as indicated in figure legends.

## Acknowledgements

Euthanized C57BL/6J mice were a gift from RR Neubig (facilitated by J Leipprandt) and the Campus Animal Resources at Michigan State University (MSU). Funding for this work was provided in part by the James and Kathleen Cornelius Endowment at MSU.

## Author contributions

Conceptualization, C.V.M. and C.H.C.; Methodology, C.V.M., S.B.G., and C.H.C.; Investigation, C.V.M., M.O.B., M.M.K. and M.H.; Writing – Original Draft, C.V.M.; Writing – Review & Editing, C.V.M., M.O.B., M.M.K., M.H., S.B.G. and C.H.C.; Funding Acquisition, C.H.C.; Resources, S.B.G. and C.H.C.; Supervision, S.B.G. and C.H.C.

